# Within-individual consistency and between-individual variation in the shapes of eggs laid by tree swallows (*Tachycineta bicolor*)

**DOI:** 10.1101/2022.02.02.478835

**Authors:** Conor C. Taff, Thomas A. Ryan, Jennifer J. Uehling, Allison S. Injaian, Maren N. Vitousek

**Affiliations:** Department of Ecology & Evolutionary Biology and Lab of Ornithology, Cornell University; Odum School of Ecology, University of Georgia

**Keywords:** egg shape, egg size, between-individual variation, scale-dependence

## Abstract

Bird eggs come in a wide variety of shapes and recent large-scale studies across species have renewed interest in understanding the factors that generate and maintain this diversity. However, these advances largely overlook the fact that egg shape also varies dramatically within species: between populations, between individuals, and between eggs produced by the same individual. We measured the shape and size of 1382 eggs produced by 205 tree swallows (*Tachycineta bicolor*) in 259 nests with many females having eggs measured in two or three years. We found that intra-specific variation in the asymmetry and ellipticity of tree swallow eggs spanned the range of typical egg shapes from 69% of species reported in a recent comparative study. Variation in shape and size was largely explained by individual identity, as the repeatability of egg shape and size was remarkably high both for eggs produced within a clutch and for clutches produced in successive years. All characteristics of mother and daughter eggs were positively correlated, although with a sample size of only 15 pairs these relationships were not significant. Female mass and structural size were correlated with the size of eggs produced, but not with their shape. Older females produced eggs that were larger, more asymmetrical, and less elliptical; this pattern was driven by a combination of both longitudinal changes in egg characteristics as females aged and by differential overwinter survival of females with particular egg characteristics. We found no evidence that any aspect of shape or size that we measured was related to nestling measurements or fate. Finally, we found that the patterns of covariation in aspects of egg size and shape between-individuals differed from that observed within-individuals, suggesting that the mechanisms driving egg shape variation might differ across these levels. A complete understanding of avian egg shapes will need to incorporate variation at multiple scales and we discuss the interpretation of our results in light of recent large-scale comparative studies that focus only on mean species egg shapes.

## INTRODUCTION

Birds vary enormously in the shape and size of the eggs they produce and explaining this remarkable variation has been a longstanding goal of ornithologists (e.g., Preston, 1969; Thompson, 1908). Recently, advances in the ease of making accurate measurements of egg shape and the ability to conduct large-scale comparative analyses have led to a renewed effort to understand what factors drive variation in egg shape between species (Duursma et al., 2018; Montgomerie et al., 2021; Nagy et al., 2019; Shatkovska et al., 2018; Stoddard et al., 2017). Despite the number of studies of egg shape, relatively few have examined variation in shape between populations, between individuals, or within individuals of the same species (but see Adamou et al., 2018; Bańbura et al., 2018).

Between species, there is general agreement that differences in life history demands and morphology contribute to egg shape variation, though the details of these patterns and their generality across different taxonomic scales is debated (Birkhead et al., 2019; Montgomerie et al., 2021; Stoddard et al., 2019, 2017). At the broadest level, much of the variation in egg shape can be explained by phylogenetic history, body mass, and egg size (Montgomerie et al., 2021; Stoddard et al., 2017). After accounting for these differences, flight ability explains additional variation in egg shape (Stoddard et al., 2017), possibly due to its association with selection on body shape and mode of locomotion (Montgomerie et al., 2021). Montgomerie et al. (2021) argue that different aspects of shape are driven by different selection pressures, with elongation being largely associated with anatomical constraints while asymmetry is an adaptation to conditions during the incubation period. More targeted studies on narrower taxonomic scales have also identified other correlates of egg shape, including pelvis shape (Shatkovska et al., 2018), climate conditions (Duursma et al., 2018), incubation behavior and chick developmental mode (Birkhead et al., 2019), nest characteristics (Nagy et al., 2019), and egg composition (Deeming, 2018), among others.

While between species variation has received considerable recent research attention, there is also widespread recognition that egg shape varies within species (Adamou et al., 2018; Bańbura et al., 2018). The causes of this variation are less clear, both because few studies measure the shape of many eggs from known individuals over multiple breeding attempts and because it is often unclear how or if explanations for egg shape variation based on between-species life history differences should apply within species or between populations. For example, Stoddard et al. (2017) found that adaptations for flight are an important driver of inter-specific variation, yet this variable is unlikely to differ much between individuals within a population (though it might between populations of a species). Similarly, Birkhead et al. (2018) argue that common murre (*Uria aalge*) eggs are shaped to provide stability on sloping ledges, but this mechanism cannot explain the maintenance of variation between individuals selected for nesting on a similar substrate.

If there is an optimal egg shape for different species (Andersson, 1978; Barta and Székely, 1997) based on differences in life history traits (e.g., Montgomerie et al., 2021; Stoddard et al., 2017), then what mechanism maintains the substantial variation in egg shape within species where life history and ecology are relatively similar between individuals? Reconciling explanations for variation in egg shape between species with the apparently large variation in egg shape that exists within at least some species requires a careful consideration of the scale over which mechanisms operate. A recent debate has focused on the general usefulness of comparing egg shape across all species of birds versus within smaller taxonomic groups that have more similar life histories (Birkhead et al., 2019; Stoddard et al., 2019). Montgomerie et al. (2021) show that relationships at the family level can differ in both strength and direction, such that broad comparisons may be subject to Simpson’s paradox. This phenomenon occurs when relationships within sub groups disappear or reverse after groups are combined (Simpson, 1951). Despite this recognition that scale is an important consideration for understanding inferences about egg shape, heterogeneity in egg shape among populations or individuals is not considered in recent comparative studies.

Because phenotypic variation is hierarchically organized (Westneat et al., 2015), the same principle of scale-specific inference can be applied to understanding variation in egg shape between populations of a species, individuals within a population, or repeated production of eggs within an individual female. Inferences derived from higher levels can always be subject to Simpson’s paradox or ecological fallacies. Recent comparative studies largely ignore the possibility of different drivers of egg shape variation at these levels by measuring only a small number of eggs per species and by conducting analyses on only a single average value for each species (e.g., Stoddard et al. 2017 and Montgomerie et al. 2021 measured a median of 8 and 3 eggs per species, respectively). There is no guarantee that relationships observed at broader levels (e.g., family level or across all birds) will explain variation between populations or individuals of the same species. A complete understanding of egg shape variation will require a scale-dependent framework (Agrawal, 2020) that makes predictions that explicitly consider the scale, based on an understanding of the mechanism(s) involved. Analyses at these different levels are complementary, rather than contradictory (Stoddard et al., 2019), but at present, very few studies provide even descriptive data on correlates of variation in egg shape within species or individuals (but see Adamou et al., 2018; Bańbura et al., 2018).

In contrast to egg shape, many studies have explored between and within individual variation in egg size, mass, or composition as it relates to maternal effects and nestling outcomes (reviewed in Christians, 2002; Groothuis et al., 2019; Krist, 2011). Christians (2002) comprehensively reviewed studies of individual variation in egg mass and found that the largest egg in a population is typically 1.5 to 2 times larger than the smallest egg. This enormous variation in egg size tends to be both highly repeatable (generally above r = 0.6) and heritable (generally h^2^ > 0.5, Christians, 2002). Some individual studies demonstrate that egg size changes with dietary supplementation (Hogstedt, 1981; Ramsay and Houston, 1997), ambient temperature (Nager and Van Noordwijk, 1992; Whittingham et al., 2007), and female age (Croxall et al., 1992; Hipfner et al., 1997), but the results of these studies are decidedly mixed and even in significant cases, the amount of variation in egg size explained is typically small (no more than 10-15%, Christians, 2002). Even female mass and size are inconsistently related to egg size variation in these studies (Christians, 2002), despite the fact that these are perhaps the most unambiguous predictors of egg size and shape between species (Montgomerie et al., 2021; Stoddard et al., 2017). While egg mass often predicts early life mass and growth of nestlings, it is less clear whether there are long term fitness consequences of initial egg mass (Christians, 2002; Potti, 1999). Given these patterns, it is possible that much of the variation in intra-specific egg shape might be explained by consistent differences in egg mass, but few studies present the data to address this question and the causes of differences in egg mass themselves remain unclear (Christians, 2002).

To begin to understand variation in egg shape at the between and within individual level, we studied variation in the shape and size of eggs produced by tree swallows (*Tachycineta bicolor*) breeding over three years. Tree swallows are a migratory cavity nesting species with a distribution spanning most of North America. They typically produce clutches containing 4-7 eggs (Winkler et al., 2020a). While we aren’t aware of prior studies directly measuring egg shape, a number of previous studies in different populations of tree swallows have documented substantial variation in egg mass (Ardia et al., 2006; Bitton et al., 2006; Wiggins, 1990) or yolk characteristics (Whittingham and Schwabl, 2002). These differences are sometimes attributed to variation in temperature, food availability, or body condition (Ardia et al., 2006; Pellerin et al., 2016; Whittingham et al., 2007), but also appear to be predominantly attributable to female identity (Wiggins, 1990).

We measured the shape of eggs of individual females over multiple years. We were initially interested in comparing the degree of variation in egg shape within tree swallows to the degree of variation described between species in comparative studies. If there is generally strong selection for an optimal egg shape matching life history traits, then we expected there would be relatively little variation in shape within tree swallows (i.e., ellipticity or asymmetry), even if there was variation in egg size associated with investment (Ardia et al., 2006; Wiggins, 1990). Next, our multi-year study allowed us to assess individual repeatability and correlates of egg shape and size both within a clutch and across multiple years. We also took advantage of the fact that some female nestlings returned to breed as adults, which allowed us to compare the shape and size of eggs produced by mothers and their daughters. Finally, we asked whether there was any evidence that variation in egg shape or size was related to nestling morphology or fate. We interpret the correlations that we observed within tree swallows in light of the results of recent comparative studies.

## METHODS

We studied tree swallows breeding near Ithaca, New York from April to July of 2019 to 2021. Tree swallows at this site have been studied continuously since 1984 and we followed well established protocols for general monitoring of breeding activity (Vitousek et al., 2018; Winkler et al., 2020b). Briefly, we checked all nest boxes every other day during the breeding season to record the timing of the onset of nest building activity, the initiation and completion of egg laying, the timing of hatching and fledging, and the fate of nestlings. This schedule also allowed us to compile accurate information on clutch size and the number of eggs that hatched at each nest, but we did not have information on the laying order of eggs within a nest.

For this study, we visited each nest during the first week of incubation and photographed eggs to measure size and shape (example photographs in Figure 1). During the years of study, many nests at these sites were included in a variety of experiments focused on manipulating environmental stressors (e.g., Injaian et al., 2021; Taff et al., 2021). However, all of these experimental manipulations began during mid-incubation, after eggs had been laid and pictures had been taken. We focus primarily on pre-treatment female and egg characteristics in this study, but we also include an analysis of nestling growth and fate from a subset of 55 nests that were not subject to any experimental manipulations (see below).

**Figure 1:**
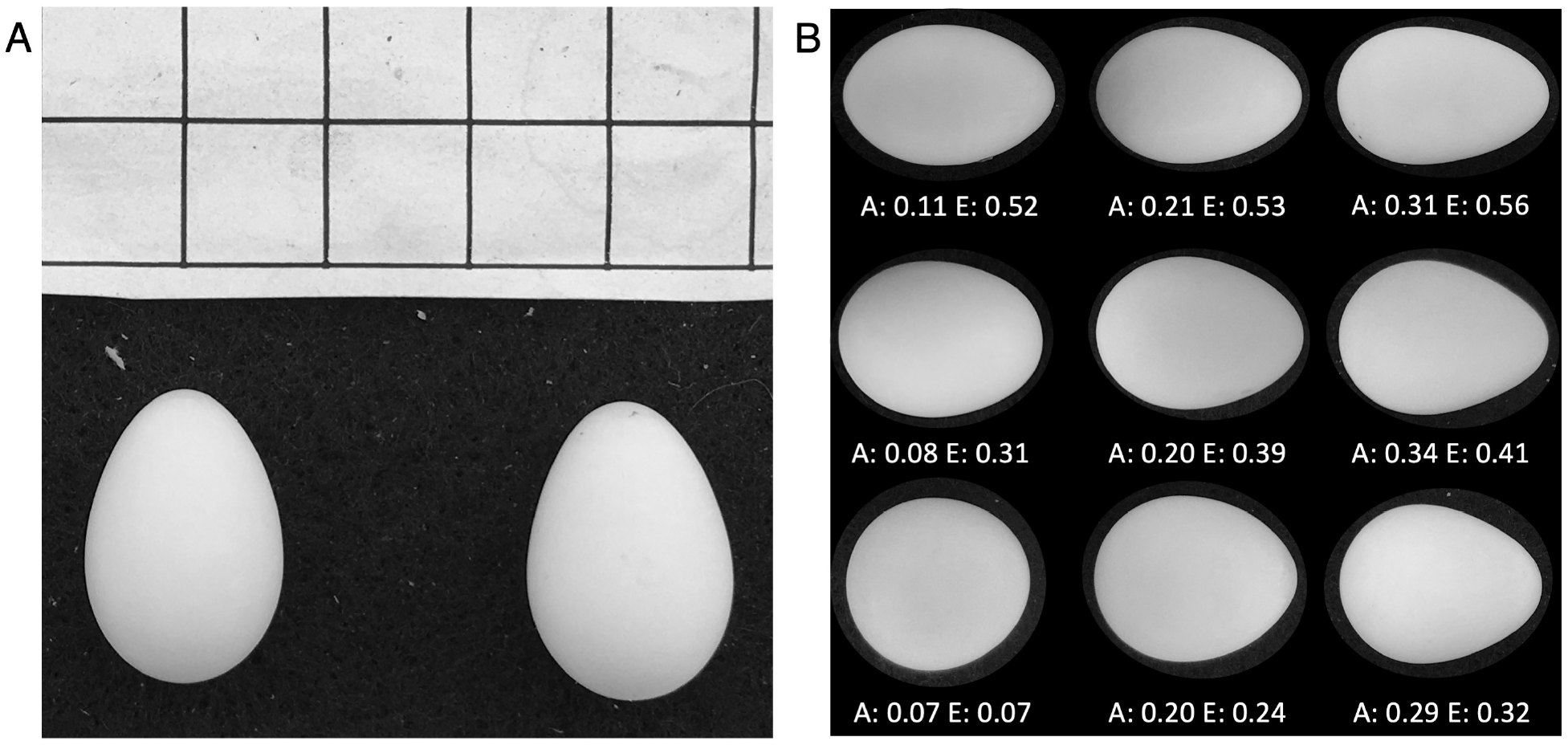
Panel A shows an example of a photograph of two eggs from the same clutch with an included scale grid for measurement. Panel B shows measurements of asymmetry and ellipticity produced from EggxTractor software with a selection of tree swallow eggs from this study. Numbers below each egg indicate the asymmetry and ellipticity. Note that eggs are not shown at scale but are rather shown at a similar size with shape maintained for illustration only.

Adult females were captured on day 6-8 of incubation between 6 and 10 am. At the time of capture, we took a series of three small blood samples and a set of standard morphological measurements that included mass, flattened wing length, and the length of the head plus bill (Vitousek et al., 2018). Any females captured for the first time were banded with a USGS aluminum leg band. Depending on the experiment, females were captured again later in incubation or provisioning and males were captured during provisioning, but those captures occurred after experimental manipulations and the data are not included in this paper.

Nestlings were banded and measured on day 12 after hatching. At this time, we measured mass, wing length, head plus bill length, and took a small blood sample. Following a return visit to experimental nests on day 15, we avoided checking nests to prevent forced fledging until day 24. We did a final nest check to determine fledging fate for each nestling.

Individuals not in the nest at this point were considered fledged and we recorded the band numbers from any dead nestlings recovered. Given our sampling strategy, we cannot link individual egg characteristics to individual nestling morphology or fates, but we can explore correlations between average egg and nestling characteristics at the nest level. While nestling characteristics and fates were recorded at every nest, we only included analyses of nests that were not subject to any experimental manipulations.

### Egg measurements

Using the photographs described above, we characterized the size and shape of eggs from each nest. To measure shape, we followed the approach developed by Baker (2002), which results in measures of the degree of ellipticity (deviation from circularity) and asymmetry (difference in shape of the two egg poles) as decribed in Stoddard et al. (2017). The measurements were performed using the `EggxTractor` software in MatLab, provided by Stoddard et al. (2017). While there are a variety of other methods to characterize egg shape that differ slightly, especially in their ability to capture highly pyriform, pear-shaped, eggs (e.g., Attard et al., 2018; Biggins et al., 2018), our approach performs well for the range of shapes covered by tree swallow eggs (Stoddard et al., 2017). To characterize the size of eggs, we used ImageJ (Schneider et al., 2012). We loaded photographs and set a scale using the scale grid that was included in every image. We then used the straight line segment tool to measure the maximum length from pole to pole (egg length) and the maximum girth (egg width) for each egg.

### Data analysis

We initially examined the overall amount of intra-specific variation in egg shape from our study in comparison with the amount of inter-specific variation presented in Stoddard et al. (2017). Given the substantial variation in egg shape between individual tree swallows, we next asked whether aspects of egg shape or size were repeatable within a female. We used linear mixed models implemented in the `rptr` package in `R` (Stoffel et al., 2017) to assess repeatability in two ways. First, we calculated overall repeatability for each egg characteristic, considering each egg measured for every female. Because eggs laid in a nest are produced under similar conditions, they may be similar to each other due to those shared conditions in addition to being similar due to intrinsic properties of the individual female. Therefore, we also calculated repeatability using average egg characteristics for each year in a subset of 59 females that were observed in multiple years. Finally, after calculating repeatability, we used a subset of 15 mother-daughter pairs that both had eggs measured to ask whether a female’s egg characteristics predicted her daughter’s egg characteristics one or more years later. Most tree swallows at our site disperse after their natal year, so the sample size for recruiting nestlings with known mothers was small (Winkler et al., 2005).

We next asked whether individual characteristics of females explained variation in egg shape or size. For these questions, we fit a series of four linear mixed models with egg shape (asymmetry, ellipticity) or size (width, length) as the response variable and with female mass, wing length, head plus bill length, and age as predictors. We included age as a categorical predictor of ‘first-time breeders’ or ‘returning breeders.’ Female tree swallows have delayed plumage maturation and first time breeders can be identified by their brown plumage regardless of prior capture history (Winkler et al., 2020a). Each of these models also included a random effect for female identity and for nest identity (to account for multiple eggs measured from the same nest). The exact age of returning breeders was not always known, but see below for a subsequent analysis comparing only known age birds. We standardized all continuous predictors to a mean of 0 and standard deviation of 1 so that effect sizes are directly comparable. Models were fit with the `lme4` package and model diagnostics were examined with the `DHARMa` package in `R` to ensure appropriate fits (Bates et al., 2015; Hartig, 2021).

After finding that female age was related to some egg characteristics, we asked whether this pattern might be best explained by a longitudinal change in egg characteristics as females age or by the selective return of females with particular egg characteristics. We used a subset of 29 females that were measured in multiple years and that were initially observed as first time breeders to ask whether egg characteristics changed longitudinally within females as they aged using paired t-tests for each egg characteristic. We used the full set of observations from 2019 and 2020 to ask whether any egg characteristics predicted the likelihood of returning to breed in the following year (we could not include 2021 females in this analysis because survival to the next year was not known). We compared the average egg characteristics in year 1 for birds that did or did not return in year 2 using t-tests. Returning the following year is not a perfect proxy for survival because adults may occasionally disperse or go undetected, but in this population previous work using the full historical database has shown that dispersal distance in adulthood is generally very small and detection of returning birds is high (Winkler et al., 2020b).

We analyzed within individual covariation in egg characteristics using a subset of 185 females that had at least 5 eggs with complete measurements. Using this subset, we calculated the pairwise correlation and R^2^ value for each pair of egg characteristics for each individual female and examined the distribution and average value of these correlations across all females. We qualitatively compared the pattern of covariation within females to that observed when using average egg characteristics between females. We examined clutch size in a set of linear mixed models including only nests that had at least 4 eggs to ask whether egg size or shape were associated with clutch size.

To ask whether any aspects of egg characteristics were associated with nestling characteristics, we used a subset of 55 nests that were not part of any experiment. For these nests, we calculated the average egg shape and size and fit simple linear models with average nestling morphology (mass, wing, head plus bill) as response variables and with egg shape and size as predictors. For nestling fate, we fit a single model with the number of nestlings that fledged and the number that did not as the binomial response and with the four egg characteristics as predictors.

All analyses and figures were produced in R version 4.0.2 (R Core Team, 2020). The complete set of code and data required to reproduce all analyses and figures is available at https://github.com/cct663/tres_egg_shape and will be permanently archived on Zenodo upon acceptance.

## RESULTS

In total, we measured the shape and size of 1382 eggs produced in 259 nests by 205 unique females. A total of 38 females had eggs measured in two years and 7 females had eggs measured in 3 years. Overall, there was enormous variation in the shape and size of tree swallow eggs. A 90% ellipse drawn based on tree swallow eggs measured in this study included the species average egg shapes for 965 out of 1400 (69%) species reported in Stoddard et al. (2017) (Figure 2).

**Figure 2:**
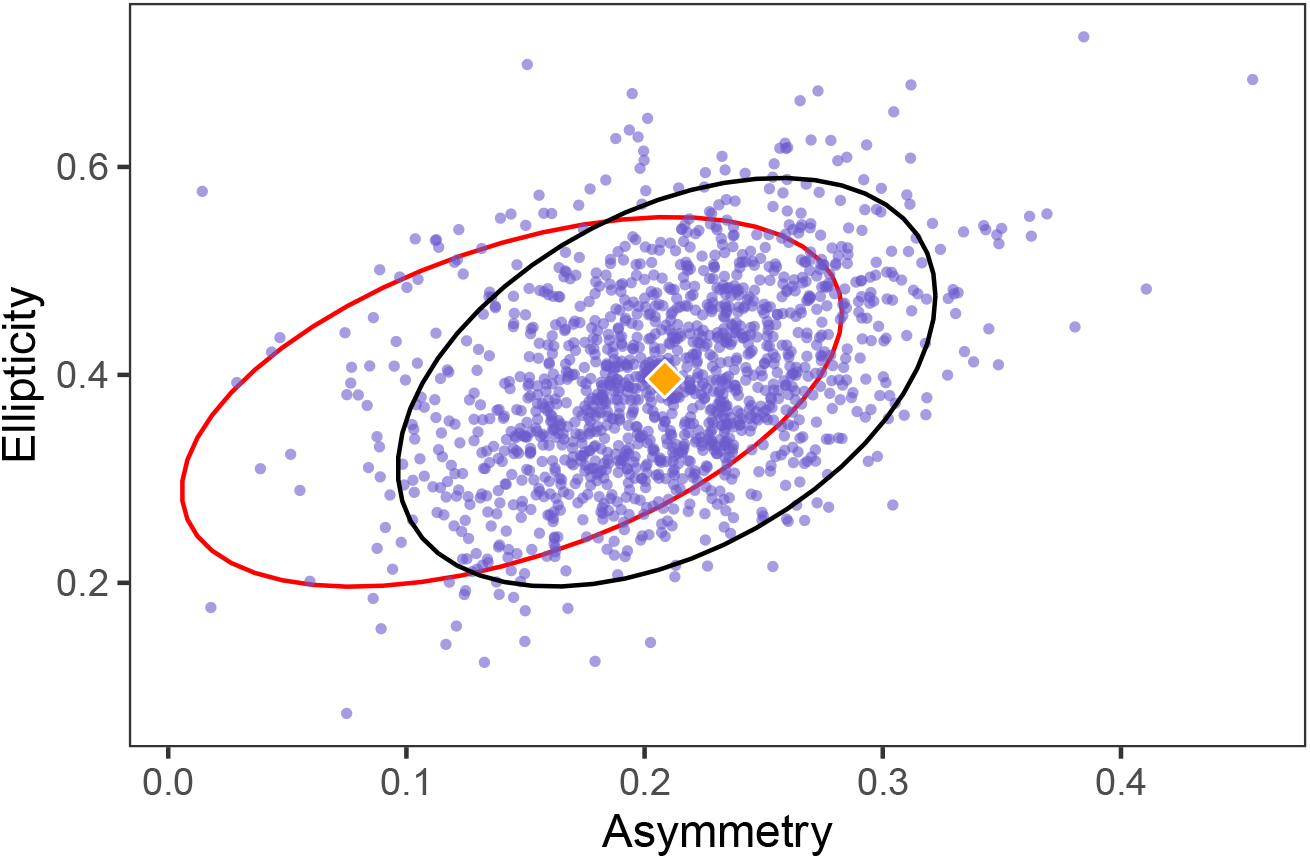
Morphospace of tree swallow egg shapes from 1382 measured eggs. Black ellipse indicates the region containing 90% of eggs. Diamond is the overall average egg shape for tree swallows from this study. The red ellipse contains 90% of the species average egg shapes from 1400 bird species included in Stoddard et al. 2017.

When comparing average egg characteristics between females, the four measures of egg size and shape that we examined were moderately to strongly correlated with each other, though each characteristic varied at least somewhat independently of the others (Table 1). However, within females there was considerable heterogeneity in the strength and even direction of the correlation between different egg characteristics for most pairs of measurements (Figure 3A). There was also little similarity between the covariation structure of egg characteristics between-females versus within-females, though the direction of the relationship was the same in all cases (Figure 3B).

**Table 1:** Between-individual correlation for each pair of egg characteristics based on the average of each characteristic for each female. Pearson correlation is shown below the diagonal and P-value above the diagonal.

|  | Asymmetry | Ellipticity | Width | Length |
| --- | --- | --- | --- | --- |
| Asymmetry |  | <0.001 | 0.74 | <0.001 |
| Ellipticity | 0.46 |  | <0.001 | <0.001 |
| Width | 0.02 | -0.45 |  | 0.004 |
| Length | 0.46 | 0.77 | 0.2 |  |

**Figure 3:**
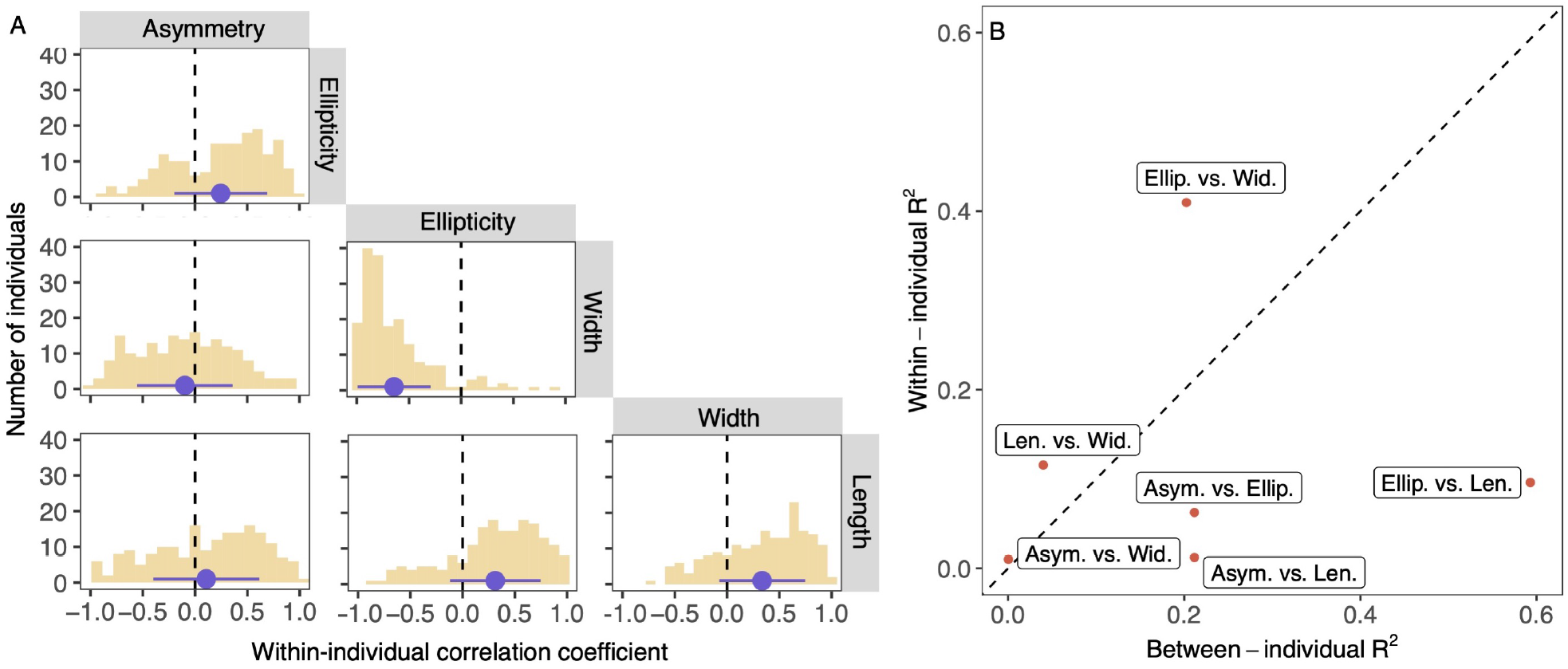
Panel A shows the distribution of Pearson correlation coefficients for within-individual correlations between each pair of egg characteristics. Dashed vertical line indicates no correlation. Blue circle is the mean and blue line denotes 1 SD above and below the mean. Panel B shows the R-squared values for the association between each pair of egg characteristics for between-individual (x-axis) and within-individual (y-axis) comparisons. Diagonal dashed line illustrates a perfect correspondence between the two axes.

### Individual repeatability in egg shape and size

Despite the enormous variation overall, individual females had remarkably high repeatability in the shape and size of eggs they produced. When considering all eggs within a clutch, both shape and size were highly repeatable (asymmetry r = 0.46, CI = 0.4 to 0.52, P < 0.001; ellipticity r = 0.54, CI = 0.49 to 0.6, P < 0.001; egg length r = 0.62, CI = 0.56 to 0.67, P < 0.001; egg width r = 0.31, CI = 0.25 to 0.36, P < 0.001). Repeatability was even higher when considering only the average egg size and shape in different years for females that returned to breed in multiple years (Figure 4 A-D; asymmetry r = 0.61, CI = 0.43 to 0.75, P < 0.001; ellipticity r = 0.65, CI = 0.5 to 0.78, P < 0.001; egg width r = 0.46, CI = 0.24 to 0.64, P < 0.001; egg length r = 0.71, CI = 0.56 to 0.8, P < 0.001).

**Figure 4:**
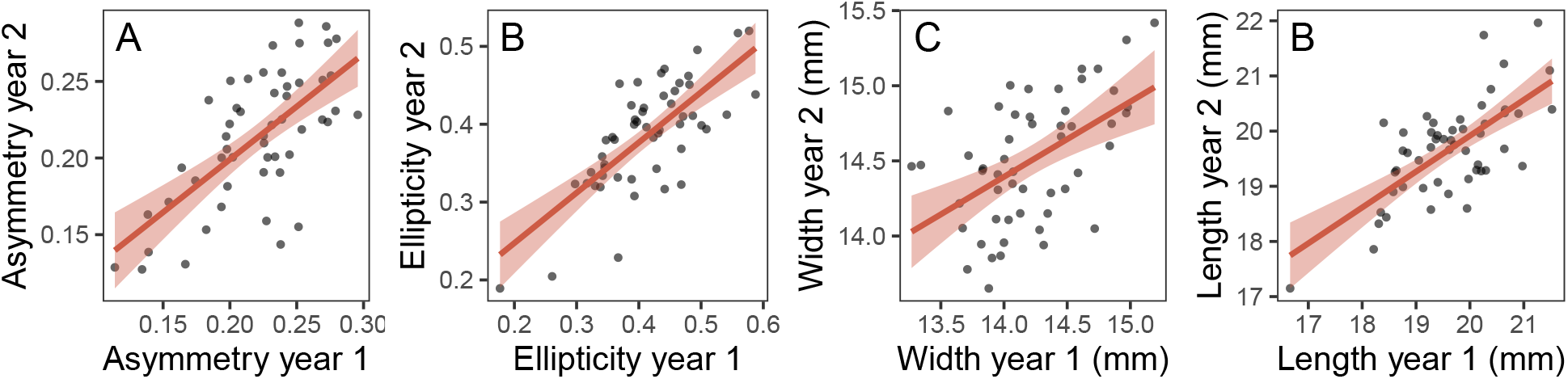
Average asymmetry (A), ellipticity (B), egg width (C), and egg length (D) for females that had egg shape measurements in consecutive years.

When comparing the eggs of mothers and their daughters, there was a positive association between all egg characteristics (Pearson’s correlation between mother and daughter for asymmetry r = 0.31; ellipticity r = 0.28; width r = 0.25; length r = 0.24). However, with a sample size of only 15 pairs, none of these relationships were significant (Pearson’s correlation test for asymmetry: t = 1.18, df = 13, P = 0.26; ellipticity: t = 1.06, df = 13, P = 0.31; width: t = 0.94, df = 13, P = 0.36; length: t = 0.89, df = 13, P = 0.39).

### Predictors of egg shape and size

In linear mixed models, no aspect of female morphology or age was related to egg asymmetry or ellipticity (Table 2). Female head plus bill length was positively correlated with egg width, while female mass was negatively correlated with egg width (Table 2; head plus bill length *β* = 0.08, P = 0.01; mass *β* = 0.04, P = 0.04). No aspect of female morphology predicted egg length, but older females produced eggs that were both wider and longer (Table 2; female age for egg width *β* = 0.24, P < 0.001; female age for egg length *β* = 0.24, P < 0.02). Despite the fact that age was significantly related to egg width and length and that morphology was significantly related to egg width, the overall amount of variation explained by the fixed effects in these models was low (Table 2; egg width full model marginal R^2^ = 0.04; egg length full model marginal R^2^ = 0.02).

**Table 2:**
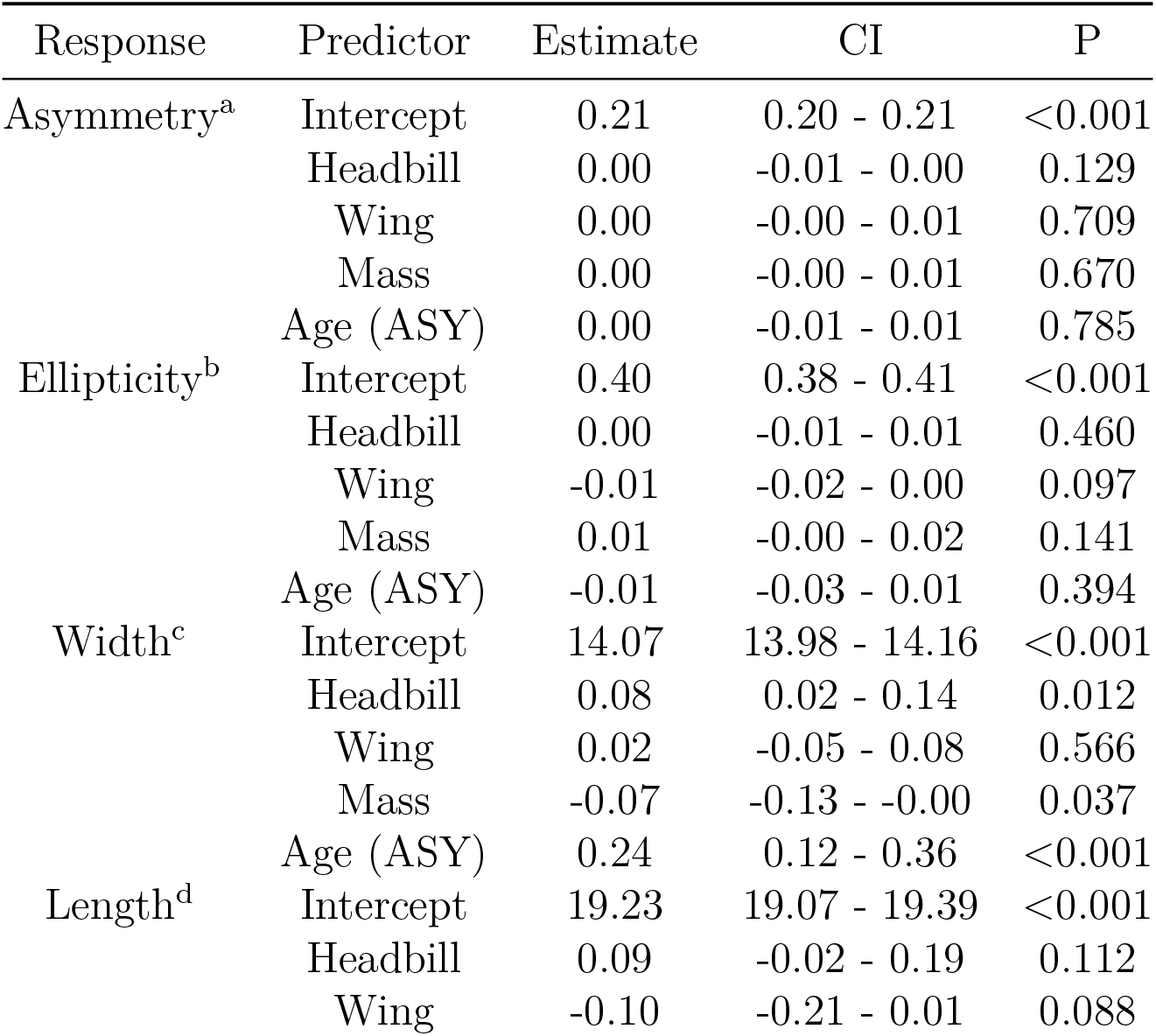
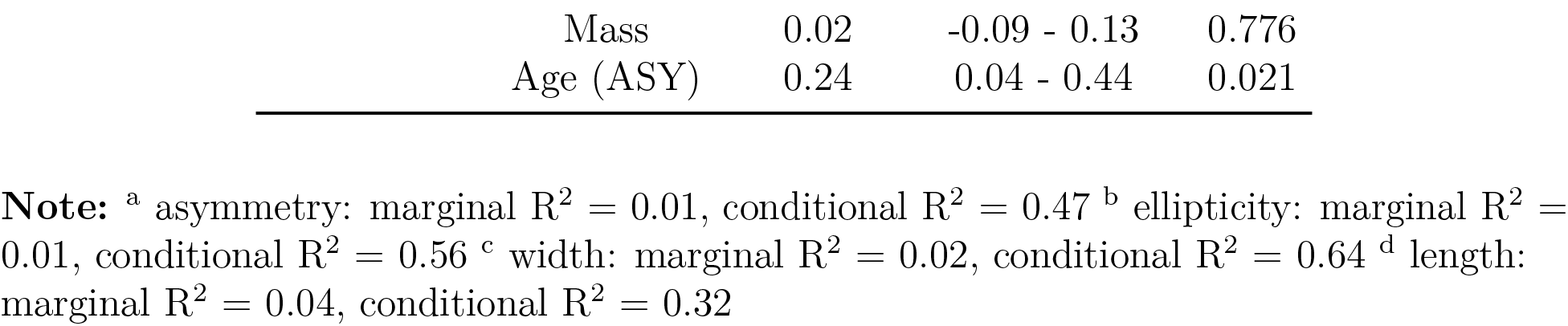
Linear mixed models with female band and nest included as random effects (*n* = 1353 eggs, 254 nests, 202 females).

Clutch size was unrelated to egg shape (asymmetry *β* = −0.0, CI = −0.01 to 0.0, P = 0.88; ellipticity *β* = −0.0, CI = −0.01 to 0.01, P = 0.59) or size (width *β* = 0.01, CI = −0.06 to 0.06, P = 0.92; length *β* = 0.07, CI = −0.03 to 0.18, P = 0.17).

### Age related changes and overwinter return

The difference in egg size with female age could arise from longitudinal increases in egg size as females age or from selective survival of females that lay larger eggs. Among females that were measured as both first time breeders and as returning breeders, asymmetry did not change longitudinally, but ellipticity decreased slightly (Figure 5 A-B; paired t-test for asymmetry: t = −0.34, df = 28, P = 0.73; ellipticity: t = −2.3, df = 28, P = 0.03). Both the width and length of eggs increased significantly from a female’s first to second breeding year (Figure 5 C-D; paired t-test for width: t = 5.31, df = 28, P < 0.001; length: t = 2.25, df = 28, P = 0.03).

**Figure 5:**
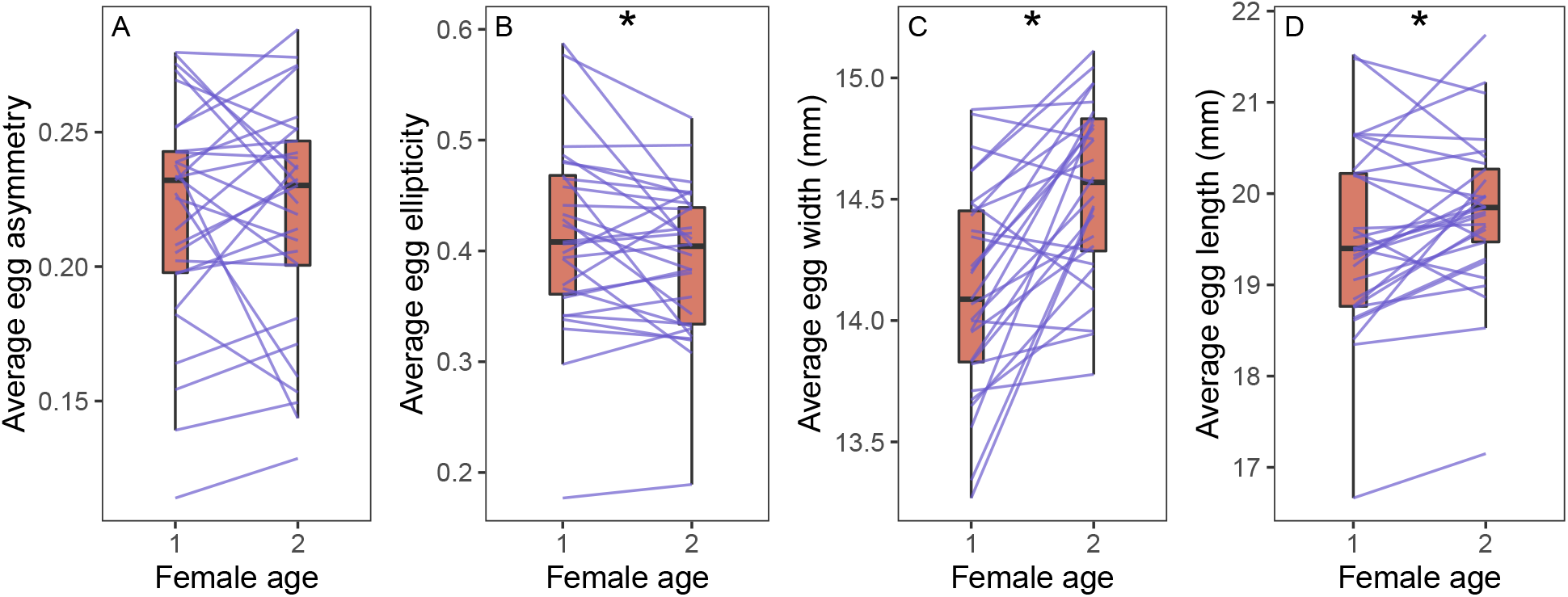
Change in average egg shape (A, B) and size (C, D) for females that were observed as one year old first time breeders and again in the subsequent year. Boxes show the mean and interquartile range for each age group. Lines connect observations from the same individual. Asterisks denote significant difference between groups.

When comparing the average egg characteristics in year 1 between females that did or did not return to our site in year 2, females with more asymmetric eggs were more likely to return the following year (Figure 6 A; Welch two sample t-test: t = 2.67, df = 152, P = 0.008). There was no difference in egg ellipticity between females that did or did not return the following year (Figure 6 B; t-test: t = 1.01, df = 160, P = 0.31). Females that laid wider and longer eggs in year 1 were more likely to return in year 2 (Figure 6 C-D; width: t = 2.60, df = 161, P = 0.01; length: t = 2.90, t = 155, P = 0.004).

**Figure 6:**
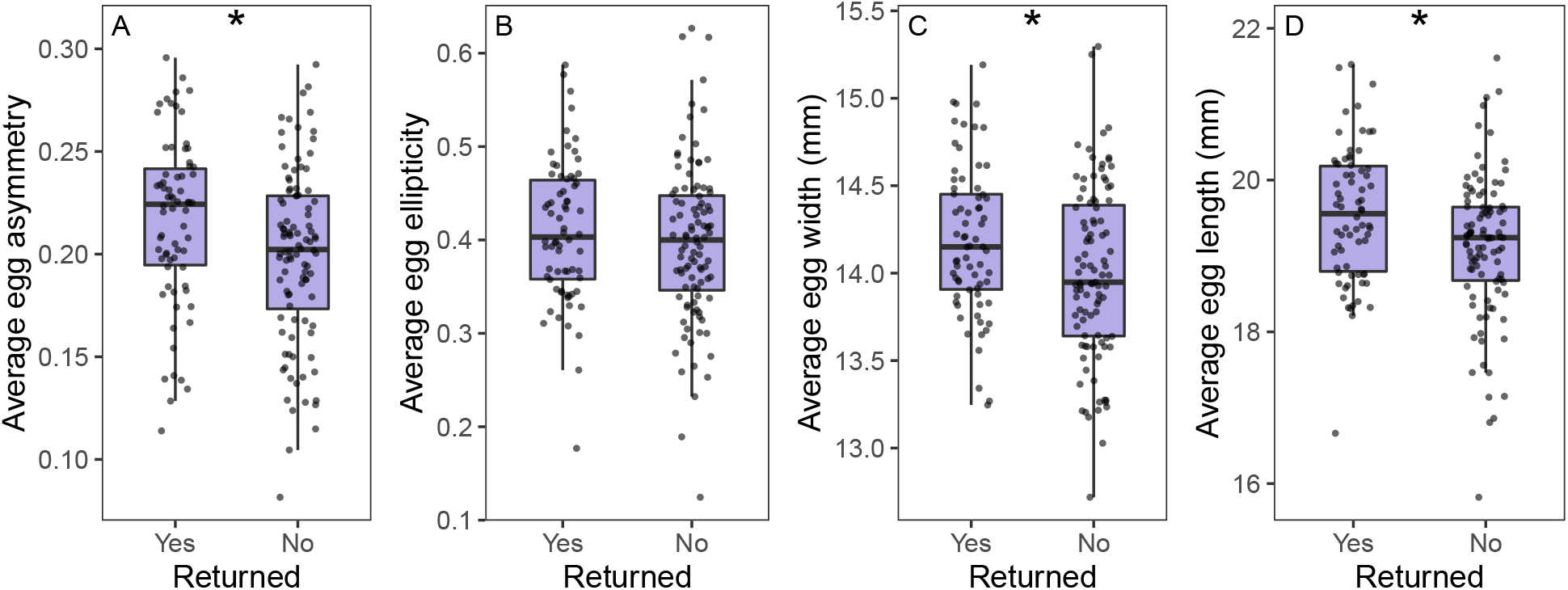
Comparison of average egg shape (A, B) and size (C, D) for females that did or did not return to breed in the following year. Boxes show the mean and interquartile range for each group. Points show average egg characteristics for each female. Asterisks denote significant differences between groups.

### Nestling characteristics and fate

There was no evidence that nestling mass or size on day 12 was related to any characteristics of egg shape or size at a nest level (linear model for nestling mass: asymmetry *β* = 0.74, P = 0.13, ellipticity *β* = 1.99, P = 0.34, width *β* = 1.55, P = 0.23, length *β* = −2.41, P = 0.20; model for head plus bill length: asymmetry *β* = 0.18, P = 0.31, ellipticity *β* = 0.55, P = 0.47, width *β* = 0.39, P = 0.41, length *β* = −0.58, P = 0.40; model for wing length: asymmetry *β* = 1.19, P = 0.29, ellipticity *β* = −0.93, P = 0.85, width *β* = −0.91, P = 0.76, length *β* = 1.13, P = 0.79). There was also no evidence that any average egg characteristic predicted the likelihood of survival to fledging among these nests (generalized linear model with binomial response: asymmetry *β* = −0.22, P = 0.60, ellipticity *β* = 1.92, P = 0.14, width *β* = −1.55, P = 0.20, length *β* = 2.17, P = 0.21).

## DISCUSSION

Surprisingly, we found that variation in the shape of eggs from a single population of tree swallows encompassed most of the variation observed in average egg shape between 1400 species (Stoddard et al., 2017). Consistent with prior studies of egg size in many species, most of this variation was attributable to between-individual differences, as we found that egg shape was highly repeatable within females and stable across multiple years. Although some aspects of egg shape and size increased as females aged and were associated with overwinter survival, we found no conclusive evidence that either egg shape or size was directly related to any aspect of reproductive performance or nestling survival. Because global patterns of egg shape diversity are, ultimately, the consequence of selection operating on variation at the within-species level, our results have important implications for understanding the mechanisms and processes that contribute to egg shape variation between species.

Perhaps the most important conclusion of our study is that explanations for variation in egg shape will likely differ subtly at each level of organization from within-individuals to between species. In a parallel to Montgomerie et al.’s (2021) suggestion that family level variation in egg shape often differs from higher order patterns, we found that the association of egg characteristics between females was quite different from that within females (Figure 3). When measuring eggs laid by the same female, there was a strong and consistent negative association between ellipticity and width, such that laying wider eggs resulted in lower ellipticity (and vice versa). While this association was present between-females, it was much weaker and instead there was a strong positive association between ellipticity and length not observed within individuals.

Between and within-individual differences in egg characteristic correlations are challenging to parse, but they suggest different mechanisms operating at different levels. Within females, there is remarkably high stability in each characteristic, so an increase in width necessarily reduces ellipticity; between-females, there is more substantial variation in both ellipticity and length, such that the same negative association is much weaker. Whether the associations between egg traits at different levels arise from pure physical constraints, resource allocation trade-offs, or selection for species level specialization is an open question and one that requires a multi-level approach (Agrawal, 2020). Inferences made from evidence at one level are not necessarily informative when applied to other levels. Explicitly defining the proposed mechanism and which levels it is expected to operate at has the potential to help reconcile apparently contradictory patterns at different levels (Birkhead et al., 2019; Stoddard et al., 2019).

Despite finding substantial variation in egg shape between females, we found little evidence that these differences were driven by overall morphology or directly related to any important fitness proxies, other than adult female survival. While a great deal is known about egg shape variation between species (Montgomerie et al., 2021; Stoddard et al., 2017), understanding what generates and maintains variation in egg shape within species is largely an open question. Given the high heritability reported for egg size in many species (Christians, 2002) and the correlations that we found between egg shape and size between-individuals, it is likely that variation in egg shape is partially genetically determined. It is worth noting that additive genetics could contribute to egg shape directly or to morphological features, such as pelvis shape (Shatkovska et al., 2018) that subsequently influence egg shape. It also seems likely that early life developmental conditions might play a strong role in determining egg size and shape throughout the lifetime. Potti (1999) found that pied flycatcher (*Ficedula hypoleuca*) egg volume as an adult was related to volume of the egg they had hatched from, but after controlling for maternal effects, early life body condition also had a strong effect on the volume of subsequent eggs and this effect persisted throughout the lifetime. This mechanism also suggests that, at least in some cases, associations between egg characteristics and female condition or performance might arise from a shared cause (early life conditions) rather than from a direct consequence of variation in initial egg shape or size.

In a meta-analysis of 283 studies that examined variation in egg size within species, Krist (2011) found that egg size is consistently related to hatching success, chick survival, growth rate, condition, and morphology. These effects are strongest early in life, but sometimes persist long after chicks have hatched (Krist, 2011). Therefore, even though we found no clear link between egg size and nestlings in our study, it is plausible that the variation in egg size that we detected could have important consequences for fitness under some conditions (e.g., when food is more limited). Prior studies of tree swallows have suggested that egg size has consequences for nestlings (Ardia et al., 2006; Bitton et al., 2006) and studies in a variety of species demonstrate that female quality or environmental conditions can generate variation in egg size (Kouwenberg et al., 2013; Kvalnes et al., 2013). This pattern might also explain the associations that we saw between egg size, survival, and age among females. Variation in individual quality is known to be an important driver of reproductive success in female tree swallows and older females are typically in better body condition and produce more offspring (Winkler et al., 2020b). In this case, the association between overwinter survival and egg size might arise because females that are higher quality or in better current condition are able to invest more in large eggs while also maintaining high survival prospects. Because of the within-individual correlations between egg shape and size, changes in overall investment (egg size) between years might have caused the smaller, but still significant, changes in shape that we detected between years. Alternatively, these correlations may be unrelated to reproductive investment *per se* and instead arise as the product of some other constraint that differs as females age, such as changes in abdominal musculature.

One question that our study cannot address is whether tree swallows have an unusually high or a typical amount of intra-specific variation in egg shape. It seems likely that species that experience strong selection for optimal egg shapes would have less between and within-individual variation in the shape of eggs produced as a consequence of consistent selection. For example, if the egg shape of the common murre is optimized to promote stability on a rocky ledge, we might expect very strong selection for the optimal shape, resulting in little variation between individuals (Birkhead et al., 2018). In contrast, tree swallows nest in cavities, produce variable clutch sizes, and have a widespread geographic distribution that results in highly variable climate conditions and resource availability during breeding (Winkler et al., 2020b; Zimmer et al., 2020). Given this life history, there may be relaxed selection on egg shape and the variation we observed may result as a by-product of selection on more consequential traits, such as morphology or yolk deposition (Ardia et al., 2006). Our study focused on only a single population, but because clutch size and environment vary predictably across the breeding range (Winkler et al., 2014; Zimmer et al., 2020), subsequent work comparing egg characteristics across different populations would be illuminating.

Large-scale comparative studies that directly address the amount of within species variation in egg shape–in addition to species mean values–also have the potential to demonstrate how selection for optimal egg shape differs across clades. Understanding which species have more or less variable egg shapes would help to clarify why the factors that explain variation in egg shape differ in their explanatory power at different scales (Montgomerie et al., 2021; Stoddard et al., 2019, 2017) and why different factors are sometimes correlated with egg shape in particular groups of birds (Birkhead et al., 2019). We suggest that comparative studies should report the values measured from individual eggs in data archives (as in Montgomerie et al., 2021) along with information on location of the original nest and female identity (i.e., if multiple eggs are measured from the same clutch photograph that should be indicated) rather than just species mean values.

In current databases, some common species have many egg measurements, however most species are represented by only a few egg measurements that might come from even fewer females from a single population (Montgomerie et al., 2021; Stoddard et al., 2017). Given the huge variation in intra-specific egg shape that we identified here, the choice of individuals in these databases could have major impacts on the results. Thus, at present these databases are not well designed to fully analyse the hierarchical nature of variation in egg shape within species, but museum collections house an estimated five million bird eggs around the world (Marini et al., 2020) and adding photographs of clutches to measure egg shape to ongoing field studies is a low cost addition. Field studies will be particularly valuable when information about the parents is also collected and when the same individuals are measured repeatedly under different conditions or after experimental manipulations.

## CONCLUSION

The past few years have seen huge advances in our understanding of how and why eggs vary in shape across all birds (Montgomerie et al., 2021; Stoddard et al., 2017). However, an understanding of how and why egg shape varies within species and within individuals has lagged behind. We demonstrate enormous variation in egg shape in tree swallows and suggest that more work is needed to understand whether this pattern is typical and the extent to which intra-specific egg shape variation itself differs between species. These studies are important for two reasons. First, there are interesting biological questions to address in their own right within species about the consequences and causes of egg shape variation. Second, understanding the mechanisms that generate variation in egg shape between individuals and how these mechanisms translate across scales of organization has the potential to clarify when large-scale comparative studies can or cannot adequately explain egg shape variation. A complete understanding of the diversity of eggs will include multi-scale explanations from individuals to all birds.

## ETHICAL NOTE

All of the procedures described were approved by the Cornell University Institutional Animal Care & Use Board (IACUC protocol 2019-0023). Captures and sampling were approved by state and federal permits to MNV.

## ACKNOWLEDGMENTS

We would like to thank the field technicians, graduate students, and postdoctoral scientists who helped with field work during the years of this study including Bashir Ali, Paige Becker, Raquel Castromonte, Navya Chamiraju, David Chang van Oordt, KaiXin Chen, Nicholas Faraco-Hadlock, Zapporah Ellis, Ginny Halterman, Jennifer Houtz, Sungmin Ko, Amanda Lazar, Alex Lee-Papastravos, Sabrina McNew, Colleen Miller, Jabril Mohammed, Natalie Morris, Yusol Park, Monique Pipkin, Olivia Rooney, Bella Somoza, and Cedric Zimmer.

## AUTHOR CONTRIBUTION STATEMENT

CCT and TAR conceived the study. CCT, TAR, JJU, and ASI collected the field data. CCT measured the photographs, analyzed data, and drafted the manuscript. MNV contributed to funding for the project. All authors provided feedback and editing on the manuscript.

## DATA AVAILABILITY STATEMENT

The complete set of code and data required to reproduce all analyses and figures is available at https://github.com/cct663/tres_egg_shape and will be permanently archived on Zenodo upon acceptance.

